# Early life stress alters lifespan trajectories of amygdala development: a cross-species model of amygdala burnout

**DOI:** 10.64898/2026.09.04.749366

**Authors:** M. Sheppard, J. Rasgado-Toledo, L.A. Trujillo Villarreal, M.C. Litwinczuk, K. L. Moran, E. McManus, R. Elliott, N.W. Duncan, E. Garza-Villarreal, Tobias Banaschewski, Gareth J. Barker, Arun L.W. Bokde, Rüdiger Brühl, Sylvane Desrivières, Herta Flor, Antoine Grigis, Andreas Heinz, Frauke Nees, Dimitri Papadopoulos Orfanos, Luise Poustka, Michael N. Smolka, Nathalie Holz, Nilakshi Vaidya, Henrik Walter, Robert Whelan, Paul Wirsching, Gunter Schumann, IMAGEN Consortium, N. Muhlert

## Abstract

Early-life stress is associated with alterations in amygdala volume, but findings vary across studies and species. Rodent models typically report increased amygdala volume, whereas human studies often find reductions or mixed effects. We propose the amygdala burnout hypothesis, which suggests that early-life stress alters amygdala developmental trajectories, producing initial volumetric increases that are followed by reductions later in life. To test this hypothesis, we examined amygdala development across rodent and human cohorts spanning adolescence to older adulthood. In rodents, chronic stress was associated with increased amygdala nuclei volumes during adolescence. Comparable increases were observed in adolescents from the IMAGEN consortium, with evidence of a dose-dependent association between childhood stress and amygdala developmental trajectories. In contrast, analyses of older adults from the UK Biobank revealed reduced bilateral amygdala volume associated with early-life stress. Together, these findings support a developmental model that reconciles previously conflicting observations across species and age groups.

## Introduction: Early life stress

Early life stress (ELS), also termed adverse childhood experiences or ACEs, encompasses a variety of stressors that may occur during childhood or adolescence, including physical, sexual, and emotional abuse; physical and emotional neglect; divorce and domestic violence^1^. A meta-analysis of 208 international studies from 1998 to 2021 estimated that 8.7% of individuals experienced three ACEs and an additional 16.1% experienced four or more^2^. Due to the high prevalence and associated negative outcomes across the lifespan, ELS constitutes a pervasive societal problem^3^. For example, individuals who experience ELS are more likely to develop both physical and mental health problems, including depression, anxiety, cardiovascular disease, and type 2 diabetes^4,5^. Furthermore, there are behavioural consequences of ELS, with a heightened likelihood of engaging in risky and dangerous behaviours, along with an increased likelihood of substance abuse problems and incarceration^1,5,6^. Consequently, ELS imposes a considerable economic burden, with an estimated annual cost of £42.8 billion in societal and health costs in England in 2020^1,7^. Despite this clear effect on behaviour and society, our understanding of the mechanisms through which ELS influences brain structure to predispose towards these negative outcomes is as yet unclear. In this work, we aim to employ a cross-species approach to understand how the brain responds to early-life stressors, with a particular focus on the amygdala, a brain region associated with threat, safety, and emotional regulation.

### The amygdala

The validity of cross-species comparisons is dependent upon the extent to which the neural structures support analogous functions across species. The amygdala is one such region with convergent evidence suggesting a conserved role in emotional learning and fear acquisition in rodents and humans^8^. In rodents, temporary inactivation via NMDA receptor blockage and protein synthesis inhibition within the central nucleus of the amygdala (CeA) can reduce the acquisition of fear learning^9,10^. Similarly, lesions to the amygdala in rodents disrupt freezing behaviour in response to foot-shock stress, impairing the rodents’ capacity for emotional learning in fear responses^11^. When the bilateral amygdalae are lesioned in humans, the perceptual ability to identify and respond to fear cues is similarly impaired^12–14^. Furthermore, using transcranial ultrasound stimulation (TUS) in humans, suppressing the amygdala, disrupted early threat learning as evidenced by slower acquisition of a conditioned threat response compared to a sham^15^ and slower learning of a threat cue compared to a safety cue^15^. In addition, a megastudy found that the amygdala was critical in early, but not late, stages of conditioning within fear learning^16^.

The amygdala also appears to be heavily involved in fear reactivity across species, namely through its cortical and subcortical connections^17^. In rodents, optogenetic stimulation paired with in vivo manipulations and ex vivo whole-cell patch-clamp recordings identified that connections between the BLA and the ventral hippocampus facilitate contextual fear and anxiogenic behavioural responses^18^. The BLA has some important projections to the striatum, thalamus and medial prefrontal cortex^19^, which are thought to be critical in regulating and producing conditioned fear responses^17^. Furthermore, damage to the CeA appears to interfere with the expression of conditioned fear responses^15^. This is considered to be due to the CeA’s projections to key brainstem nuclei such as the periaqueductal grey, parabrachial nuclei and the solitary nucleus, which control visceral and behavioural responses to fear and stress^17,20^. Comparatively, fMRI studies in humans have shown that exposure to fear-inducing stimuli has been associated with amygdala activation^21,22^. A meta-analysis of 55 PET and fMRI studies highlighted that the amygdala is consistently associated with fear and emotional responses^23^. Together, these findings highlight the conserved role of the amygdala in fear and threat processing across species, supporting its use as a translational target for cross-species investigations of ELS.

### Volume change in the amygdala associated with ELS

MRI-derived volume change in the amygdala has been documented following ELS; however, the direction and scale of changes have not always been consistent across rodent and human models. There are several potential reasons as to why we see these inconsistencies. Firstly, the directionality of effects may be specific to the type of stressor experienced. For example, institutionalised children, who are likely to experience neglect and abuse^24^, demonstrate larger amygdala volumes than non-institutionalised counterparts^24,25^, with these volume changes linking to deficits in emotion regulation^24^. However, in non-institutionalised samples who also experience neglect and abuse, reduced volume was identified^26^. Additionally, a review on neurobiological changes associated with childhood maltreatment identified 8 studies reporting significant reductions in amygdala volume, 13 that claimed no differences, and 4 that reported amygdala volume increases^27^. Furthermore, latent class modelling demonstrated no association between types of ELS and amygdala volume change^28^. Therefore, although stressor type may contribute to heterogeneity across studies, the mixed pattern of findings across different forms of adversity suggests that stressor type alone is unlikely to fully explain the inconsistencies in the literature.

Alternatively, the inconsistencies may reflect effects of neurodevelopmental processes. The trajectory of amygdala growth in non-stressed samples typically shows increases in size and volume by 40% between the ages of 8-18 years^29,30^, with 30% of this growth being attributed to the maturation of neurons within the BLA^31^. However, the growth between childhood and adulthood is non-linear^32^; therefore, after reaching peak volume, amygdala volume appears to stabilise or decline across later development and ageing. As only one static measure of volume is often considered in human models at different developmental points across studies (i.e., childhood versus later life), this could result in conflicting and contradictory findings.

Inconsistencies seen in cross-species comparisons may then be a consequence of incompatible assessment points. For example, human studies often image retrospective adult samples who previously experienced ELS, whereas rodent models assess volume changes during childhood and/or adolescence, either whilst the stressor is still occurring or immediately after cessation. Consequently, the effects of ELS within rodent models may be more consistent with an increase in volume often being identified^33,34^; however, human findings are more inconsistent and often do not align with the animal models. Thus, apparent cross-species inconsistency may partly reflect a mismatch in developmental sampling rather than a true divergence in biology. This could help explain the discrepancy in findings between rodent and human studies of ELS.

Furthermore, a meta-analysis assessing volume changes in the amygdala associated with ELS identified a non-linear U-shaped relationship when considering the age of the sample and the direction of the effect^35^. The findings suggest that in ELS-exposed populations, younger samples exhibit increases in amygdala volume, whereas older participants show volume decreases^35^. This is an important distinction, as if the amygdala appears to mature differentially for individuals who experienced ELS compared to those who did not, the life stage at which the volume is assessed could be key to distinguishing why findings between rodent and human models diverge. This may, therefore, offer a sensible explanation for both the discrepancy within human findings and cross-species comparisons.

Although the rodent and human models examined here capture different forms of developmental stress exposure, all involve prolonged activation of stress-responsive neurobiological systems and therefore provide complementary translational models of adversity. Cross-species comparisons of stress-related amygdala development are often complicated by methodological differences between animal and human studies, including the use of histology in rodents and MRI-derived volume measures in humans41. To reduce this source of heterogeneity, we used MR-based measures of amygdala volume across species. We first tested whether developmental chronic restraint stress was associated with altered amygdala volume in rats. We then examined whether early stress was associated with altered longitudinal trajectories of amygdala volume from adolescence into early adulthood in the IMAGEN cohort. Finally, we tested whether childhood stress was associated with lower amygdala volume in mid-to-later adulthood using the UK Biobank. Based on this, we postulate a conceptual ‘amygdala burnout model’ which suggests there are trajectory differences in amygdala development for those who experienced ELS compared to those who did not, with ELS being associated with steeper amygdala growth trajectories in the short term and lower volume in the long term (Fig 1). We hypothesised that ELS would be associated with increased amygdala volume or steeper amygdala growth during adolescence, but reduced amygdala volume in later adulthood.

**Figure 1.**
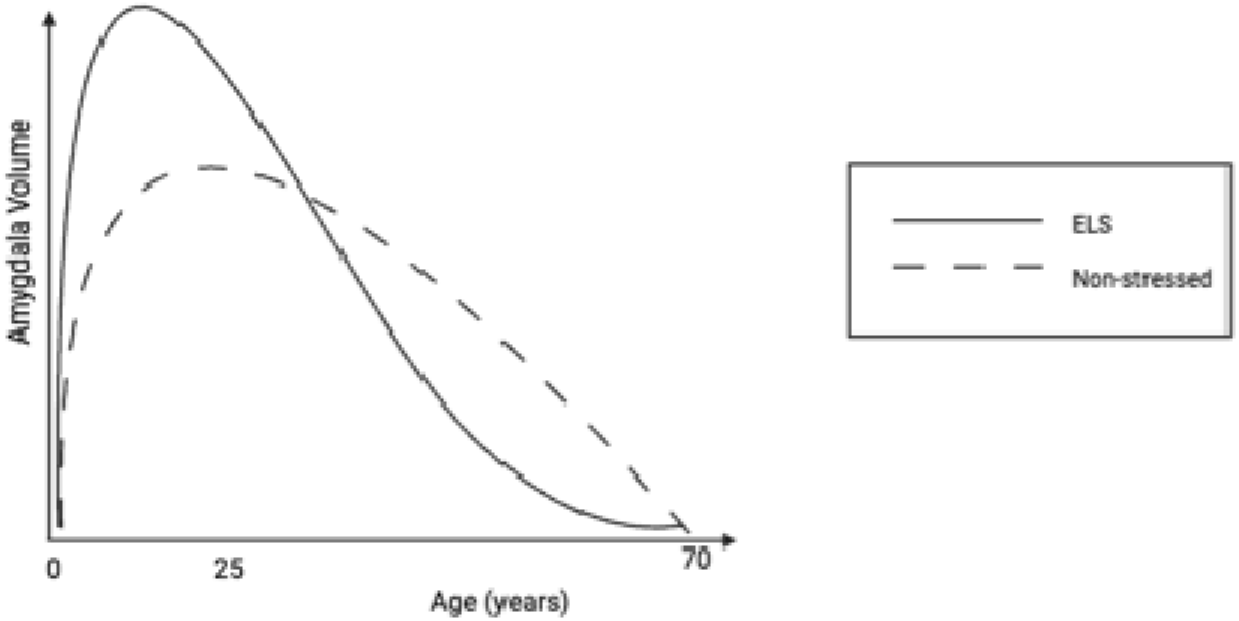
The proposed ‘amygdala burnout’ model. This predicts that amygdala volume increases in the short term, early in life, following significant stressors, but then plateaus or decreases in later life.

## Findings

### Rodent Models

Chronic restraint stress was associated with significantly greater age-related amygdala volumetric trajectories compared with controls (t = 2.02, DF = 874.07, q < .05). Voxel-wise analyses localised these effects to the bilateral basolateral amygdala (BLA) and left central amygdala (CeA). Peak effects were identified within the left CeA (peak voxel coordinates = −2.87, 0.41, 0.08; β = 0.00149, SE = 0.00012, t = 12.09, p < .001) and bilateral BLA (peak voxel coordinates = 4.84, −0.08, −0.57; β = 0.0014, SE = 0.0002, t = 5.99, p < .001) (Fig. 2).

**Figure 2.**
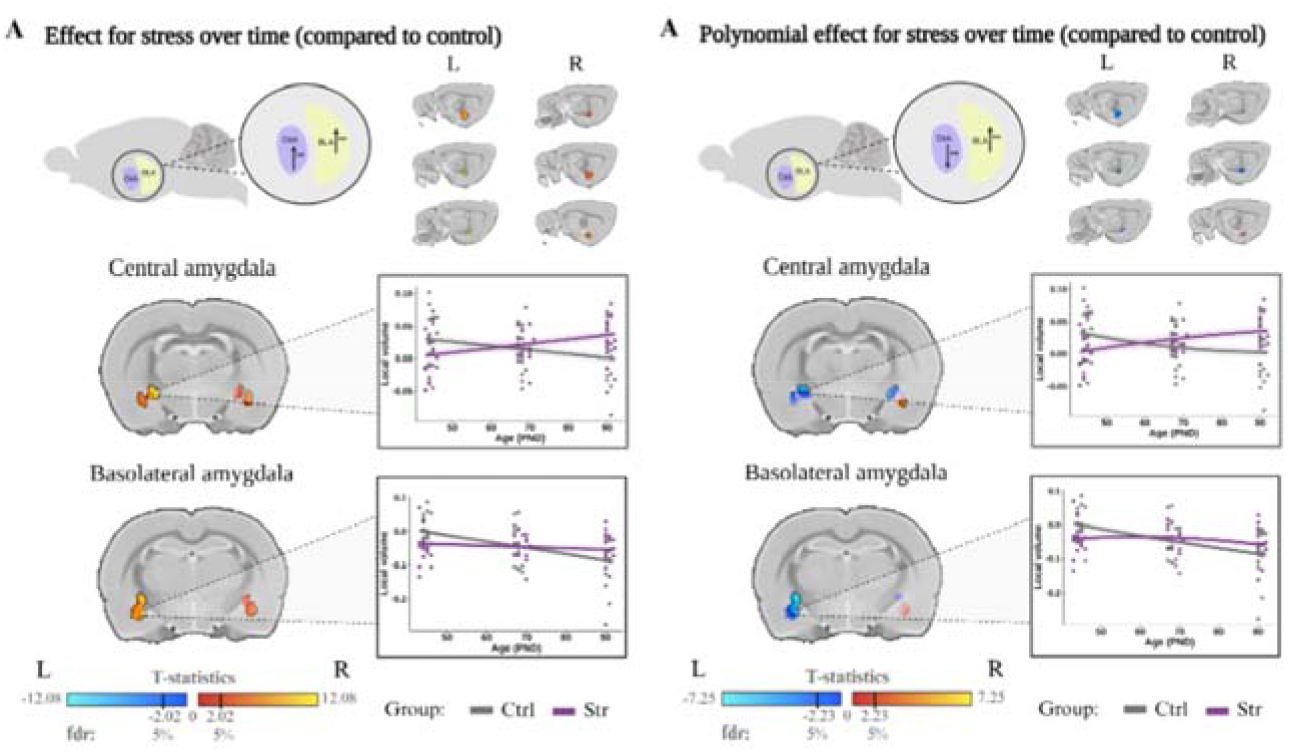
Structural differences in the amygdala between chronic restraint stress and control groups were examined using voxel-wise volumetric trajectories over time. A) Coronal slices of a rat brain average with statistically significant volumetric differences displaying bilateral volume increases in the basolateral nucleus (BLA) and left central nucleus (CeA) of the amygdala in rodents. The right panel shows the plot of relative volume change (mm3) over the three points for a peak voxel in the right BLA and in the left CeA. Colour map denotes t-statistic map of group volumetric effects thresholded at False Discovery Rate (FDR) 5%. B) Polynomial (quadratic Age² × Group) models examined non-linear volumetric trajectories over time. A) Age² × Group analysis revealed attenuated stress-related effects localised predominantly to the bilateral basolateral nucleus (BLA) and left central nucleus (CeA) of the amygdala, with reduced effect magnitudes compared to linear models. Right panel plots relative volume change (mm³) across age spans for a representative peak voxel, illustrating the quadratic trajectory.

To examine whether non-linear (quadratic) age trajectories better captured stress-related amygdala changes, we tested polynomial models incorporating Age² terms. Polynomial models (Age² × Group) revealed attenuated effects compared to linear models (FDR threshold: t = 2.231, DF = 871.46, q < .05), with overall stress-related volumetric effects that were less robust across the amygdala. Peak effects in the polynomial model remained localised to similar anatomical regions (right BLA: t = 5.352; left CeA: t = −7.251; right CeA: t = −6.697), though some effects showed directional reversals compared to the linear model, suggesting that age-related trajectories are better characterised by linear rather than quadratic functions. Exploratory analyses identified sex-specific effects in the rodent sample. As sex differences were not a primary aim of the study and were not replicated in either human cohort, these analyses are reported in the Supplementary Information (Supplementary Note 1).

### Adolescent Sample

To assess whether childhood stress influenced amygdala volume, a linear mixed-effects model was fitted. A significant age x CTQ interaction (β = 2.38, SE = 0.96, t = 2.49, p = .013) was identified, indicating that greater childhood trauma was associated with steeper increases in amygdala volume across adolescence and early adulthood (Fig 3). To account for inter-individual variability in developmental trajectories, subject-specific random age slopes were fitted. A likelihood-ratio test demonstrated that models including random age slopes fit significantly better than random-intercept-only models (χ²(2) = 342.7, p < .001). Results were consistent across both model specifications (Supplementary Note 6; Supplementary Table 1).

**Figure 3.**
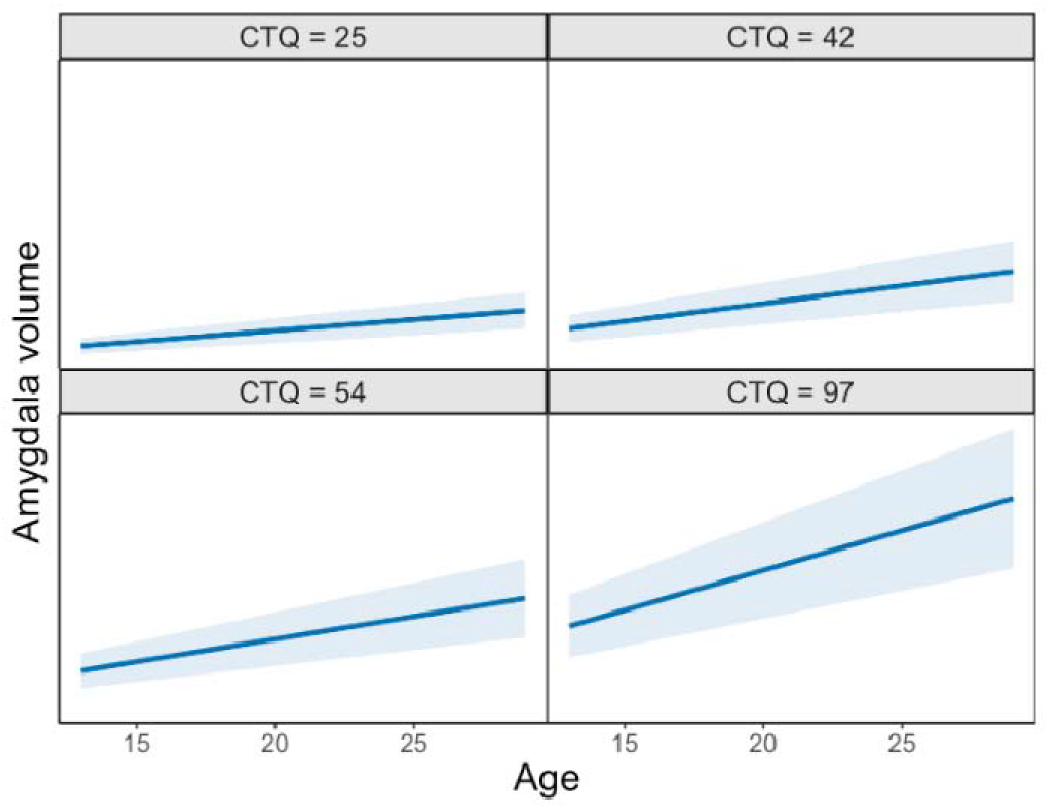
Modelled developmental trajectories of amygdala volume across adolescence and early adulthood depending upon the severity of the cumulative stress experienced. Trajectories shown span the observed range of the sample (CTQ=25-97). Shaded ribbons represent 95% confidence intervals.

**Figure 4.**
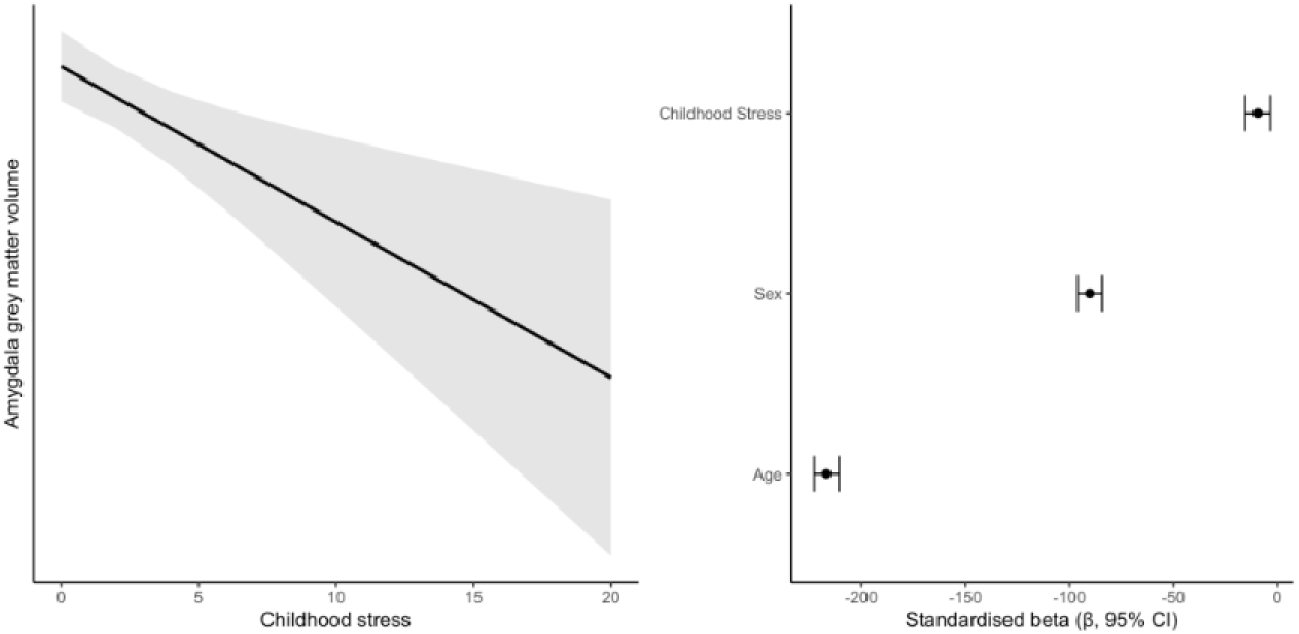
Higher levels of childhood stress are associated with lower amygdala volume in mid to late adulthood. (A) Adjusted relationships between ELS and amygdala volume when controlling for age and sex.The ribbon represents 95% confidence intervals. (B) Standardised regression coefficients (β ± 95% CI) illustrating the effect sizes of childhood stress, age and sex.

To examine whether childhood stress influenced amygdala development non-linearly, a polynomial mixed-effects model including both linear and quadratic age terms was fitted. Although a significant quadratic effect of age was observed (β = -2.12, SE = 0.82, t = -2.58, p = .010), neither the Age × CTQ interaction (β = -0.54, SE = 0.96, t = -0.56, p = .576) nor the Age² × CTQ interaction (β = -1.53, SE = 1.14, t = -1.34, p = .180) reached statistical significance. These findings do not support a non-linear influence of childhood trauma on amygdala developmental trajectories across adolescence and early adulthood. Exploratory analyses did not provide evidence that the association between early life stress and amygdala developmental trajectories differed by sex (Supplementary Note 2).

### Older Adult Sample

Linear regression analyses indicated that higher levels of ELS were associated with reduced amygdala volume in later adulthood after adjusting for age and sex (standardised β = −0.018, t(25,292) = −3.16, p = .0016). The overall model was significant (R² = 0.20, F(3, 25,292) = 2161, p < .001).

A follow-up voxel-wise analysis within the bilateral amygdala ROI identified significant localised reductions in grey matter volume in participants with high levels of ELS (p < .05, small-volume corrected). Peak effects were observed in the right amygdala (MNI coordinates: x = 18, y= -9, z = -14) and left amygdala (x= -18, y=-10, z = -14) (Fig 5). No evidence was identified for sex-specific associations between childhood adversity and amygdala volume in the UK Biobank cohort (Supplementary Notes 3 and 4).

**Figure 5.**
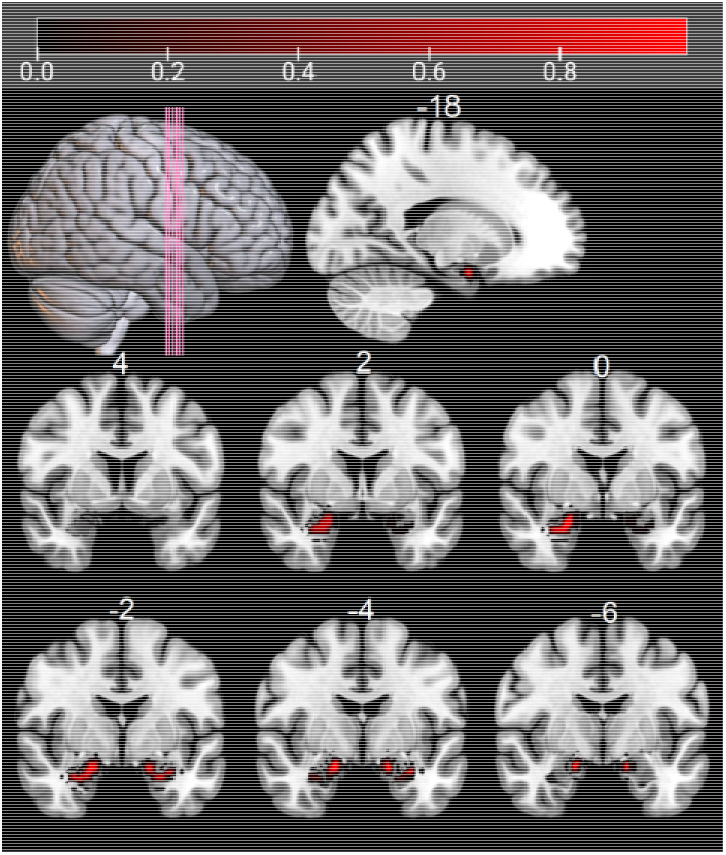
Voxel-wise localisation of ELS-related reductions in amygdala grey matter volume in mid to later adulthood. Significant voxels are shown following small-volume correction within the bilateral amygdala mask (p <.05 SVC-corrected). Numbers indicate MNI slice coordinates.

## Discussion

In this study, we used a cross-species approach to assess amygdala volume change associated with ELS. We tested the amygdala burnout hypothesis, which postulates trajectory changes in amygdala volume development for those who experience ELS. Namely, we hypothesised that in the short-term, there would be steeper developmental trajectories in the amygdala, whereas in later life, the amygdala volume would decrease compared to controls. Our findings suggest nuclei-specific volume increases in the BLA and CeA in stressed adolescent rodents and altered amygdala developmental trajectories in human adolescents with high CTQ scores. The findings observed in our adolescent human sample align closely with the juvenile rodent sample. This is particularly important as the age range represented in the adolescent sample is analogous to the developmental stage represented by the rodents^36^, suggesting that in both cases, similar processes may occur. Furthermore, our findings suggest reduced amygdala volumes in mid- to later-life adults who had experienced high childhood stress. This finding is consistent with the later-life component of the amygdala burnout hypothesis. These findings converge in suggesting that differences in developmental trajectories could help explain discrepancies in previous cross-species work, highlighting the importance of lifespan stage in assessing the impacts of ELS on amygdala volume.

The short-term nuclei-specific volume increase identified in our rodent models is in line with previous rodent work^33,34^. This has shown BLA volume increases following stress, which at histology was linked to increases in dendritic arborisation of pyramidal and stellate neurons^34^ and increases in spinogenesis and spine density in BLA spiny neurons^33^. The BLA is of particular importance as 30% of the amygdala’s normative development is driven by the maturation of BLA neurons^31^. This highlights a potential developmental vulnerability: if this region is particularly sensitive to ELS, stress-related alterations of these maturation processes could disproportionately drive early increases in amygdala volume. Furthermore, as disinhibition of the BLA is required to adaptively respond to stress^17^, volume changes in this region following frequent stress-induced activation could be particularly marked.

Volume increases in the CeA associated with stress were also identified in our rodent models. Activation studies have demonstrated that stressors in childhood upregulate corticotropin-releasing hormone (CRH) expressing neurons in the CeA and that this is associated with increased anxiogenic behaviours^37–39^. As the CeA is an output nucleus within the amygdalar complex, it is thought to regulate anxiety responses through downstream GABAergic projections, increasing further CRH release^40^. CRH expression in the CeA is an important adaptation to chronic stress responsivity, with increased synthesis associated with dysregulated HPA responses via increased arginine vasopressin concentration in the paraventricular nucleus of the hypothalamus and reduced glucocorticoid negative feedback^41^. Transient psychological stressors have been demonstrated to increase CRH transcription and associated mRNA levels, which soon return to baseline after exposure^42^. However, chronic and repeated stressors, such as systematic immobilisation stress, increase CRH transcription and elevations in mRNA cumulatively after each stress exposure^43^. As such, the volume change identified in the CeA within this study may be explained by sustained stress-induced adaptations involving CRH signalling. Such adaptations may represent one mechanism through which chronic stress alters amygdala structure during sensitive developmental periods.

Whilst we provide longitudinal evidence for altered amygdala volume trajectories in adolescence, we only have cross-sectional data in the mid-to-later life sample. As such, we cannot yet test the trajectory of the downward slope in our model. However, as we have identified that amygdala volumes in mid-to-later life adulthood are smaller in those who experienced ELS, we can postulate three potential trajectory models (Fig 6). Firstly, the decline in volume could be quite steep and mirror the accelerated developmental trajectory identified in adolescence. Secondly, the trajectory could begin to fall steeply and then converge with normative development. Finally, after the amygdala volume peaks, the trajectory of the stressed sample could match the normative trajectory decline. Without having longitudinal data for mid-to-late adulthood, we cannot ascertain which of these models best explains the lifespan trajectory of development. Therefore, longitudinal studies of human amygdala volume change in older adults associated with ELS should be conducted to allow us to appropriately map lifespan development and provide further clarity on the discrepancy in previous findings.

**Figure 6.**
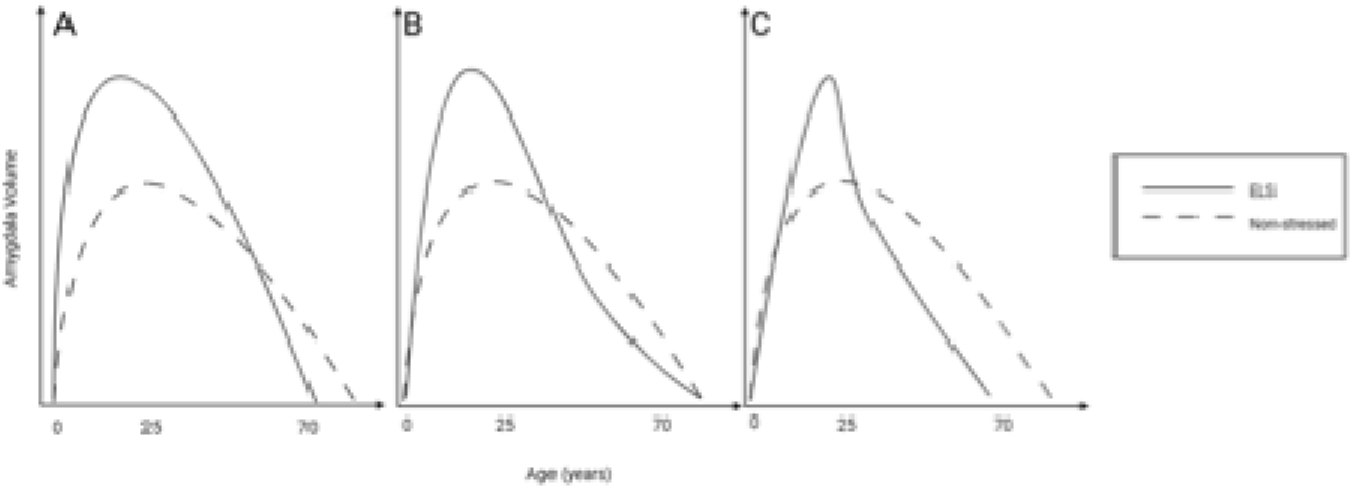
Three potential models of amygdala burnout. Our studies demonstrate altered trajectories during adolescence, but without longitudinal data, we hypothesise that in mid to older adulthood, it could either (A) have a steeper decline, (B) converge to return to the same final end-point volume or (C) maintain the same trajectory as individuals without ELS exposure after peak burnout.

Recent lifespan mapping studies have identified distinct developmental epochs, with adolescence extending into early adulthood and substantial reorganisation of brain network topology occurring around the transition to adulthood)^44^. This framework is broadly compatible with the amygdala burnout hypothesis. Our findings suggest that ELS-associated increases in amygdala volume or steeper developmental trajectories are present throughout adolescence, whilst previous work has reported similar associations up to at least 28 years of age^44^. In contrast, our UK Biobank findings indicate reduced amygdala volume in mid-to-later adulthood, consistent with the possibility that the divergence between ELS-exposed and non-exposed individuals emerges during early adulthood^45^. Although the precise timing of this ‘point of burnout’ remains unknown, these findings provide a tentative timeframe over which peak amygdala volume may be reached before subsequent reductions in volume compared to normative development.

When considering how ELS influences brain structure across the lifespan, it would be remiss to ignore the potential confounding effects of additional adulthood stress. Those who experience adversity in childhood are more likely to experience significant stress, such as sexual abuse, later in life^46^. The effects of adulthood stress on the volume of the amygdala are currently unclear. Cross-sectional work has identified that in humans, experiences of adulthood stress alone, or in combination with ELS, do not influence the structure of the amygdala in terms of both macrostructure and microstructure^47^. However, rodent models of ‘double hit’ stressors that induced stress in childhood through a maternal separation paradigm and a foot shock stressor in adulthood, identified increases in volumes compared to control animals after both childhood and subsequent adulthood stressors^48^. One interpretation is that this increase represents a transient adaptive response to stress. However, the impact of this second significant stress may alter the trajectory of amygdala volume further. Therefore, future research should investigate how stressors at different time points in the lifespan interact and influence amygdala volumes.

Our multi-species and multimodal work are consistent with a lifespan amygdala burnout model in which ELS is associated with larger amygdala volumes and altered developmental trajectories during adolescence, but lower amygdala volume in later adulthood. These results provide evidence compatible with the developmental account of the inverted U-shaped relationship between amygdala volume change and ELS proposed in previous meta-analytic work^35^. These findings highlight the importance of developmental timing when assessing neurobiological consequences of ELS and provide a framework for future longitudinal studies of stress-related brain development across the lifespan.

## Methods

### 1. Rat Adolescent model

First, we studied longitudinal changes in amygdala volume following adolescent chronic restraint stress.

#### Animals

Wistar Rats (n=34, females=16, acquired in two batches) were obtained at postnatal day 21 (P21) from the Institute of Neurobiology’s vivarium from an ethanol and stress study^49^ and randomly divided into groups. For this study, we used a sub-sample of 2 groups (no ethanol): Stress (n=17, females=8) and Control (n=17, females=8). The rats were housed in same-sex rooms maintained at a temperature of 26°C and a humidity level of 60%, with 2 cm of wood chip bedding, under a 12-hour light/12-hour dark inverted cycle (lights on from 19:00 to 07:00).

All experimental procedures and animal care were conducted in accordance with the “Reglamento de la Ley General de Salud en Materia de Investigación para la Salud” (Health General Law on Health Research Regulation) of the Mexican Health Ministry, adhering to the “Guide for the Care and Use of Laboratory Animals” and the “Norma Oficial Mexicana” (NOM-062-ZOO-1999). The animal research protocols were approved by the Ethics Committee of the Neurobiology Institute of the National Autonomous University of Mexico under project number A113.

#### Procedure

The chronic restraint stress protocol was conducted starting at postnatal day 45 to induce stress following established protocols^50^. All rats were weighed daily at the start of the 12-hour dark cycle to measure the weight gain change over age, as a measure of stress-induced response^49,50^. Daily weight measurements were collected as part of the validation of the chronic restraint stress paradigm. The behavioural and physiological effects of the stress manipulation, including altered weight gain trajectories, have been reported previously in the parent study^49^. Restriction was applied intermittently for three hours (starting at 11:00 AM), five random days per week, to minimise habituation. The restrictor was made of an acrylic tube and featured a double adjustment to accommodate the rat’s size.

Neuroimaging data were acquired at the National Laboratory for Magnetic Resonance Imaging (LANIREM) using a 7 Tesla Magnetic Resonance Imaging (MRI) scanner (Bruker Pharmascan 70/16 US) with a 2 x 2 array surface rat head coil, through Paravision v.7.0 interface software. The sequences included high-resolution T2-weighted images (sMRI), resting state functional MRI (rsfMRI) sequences, and diffusion-weighted images (dMRI) with a total scan time of 54 minutes and 54 seconds. Here, we only used the sMRI images, acquired with a 3D FLASH sequence T2w with 2 repetitions, TR = 30.76 ms, TE = 5 ms, flip angle = 10°, FOV = 25.6 x 19.098 x 25.6 mm, and an isometric voxel of 160 microns. Three time points (T1, T2 and T3) were used in the present study. After a baseline MRI acquisition (P43 ± 1 day), a second scan was performed at P67 ± 2 days, with a final scan at P91 ± 1 day.

For the acquisition, all rats were weighed and anaesthetised outside the scanner. Unconsciousness was achieved by inducing anaesthesia with vaporised isoflurane (∼4%) in a transparent polypropylene chamber. Once anaesthetised, the rats were placed in a supine position at the scanner’s entry. Anaesthesia was maintained using a 50/50 mixture of isoflurane and oxygen, administered through airflow to the nose at a concentration of 0.6 to 1% isoflurane. Cardiac and respiratory rates were continuously monitored to detect waking during the MRI acquisition.

#### Magnetic Resonance Imaging analysis

Two consecutive structural T2-weighted 3D images were acquired at each of the three sessions (T1, T2, T3). The two images were averaged to enhance the signal-to-noise ratio (SNR). We then conducted a thorough visual inspection of the average scans to identify any low-intensity scans or artefacts that could affect data quality for preprocessing for Deformation-Based Morphometry (DBM) analysis. The pre-processing stage included intensity and inhomogeneity normalisation, centring of the image, and denoising by using an in-house pipeline built on MINC-toolkit-v2 and the Advanced Normalisation Template (ANTs) tools^51,52^. In this study, a non-linear approach was utilised to average each scan between sessions, thereby creating a subject template. Subsequently, an iterative process was employed to average each subject template, resulting in the generation of an overall template that encompassed all subjects^53,54^. The final preprocessing step entailed the iterative normalisation of scans from native space to template space, the calculation of log-transformed Jacobian determinants for each subject and session, and the implementation of a final smoothing step employing a 1mm kernel. The DBM analysis was delimited by employing a binary mask of the basolateral (BLA) and central amygdala (CeA) regions. This mask was manually created using the Paxinos atlas^55^ as a reference on the created average template. It was then used as an ROI mask for statistical analysis. The final results of this process were deformation maps for each session per subject to measure the local volume change along the protocol.

#### Statistical Analysis

All statistical analyses were conducted using R (Version 4.2) in RStudio (version 2024.04.1 Build 748). Before the analysis, sensitivity tests were conducted to remove any outliers from the dataset using interquartile ranges (IQR). Outliers were defined as values that exceeded three times the IQR and were removed from the dataset. Age was mean centred prior to analysis. To assess amygdala volume change over time associated with ELS, a 2 linear mixed-effects model was fitted:

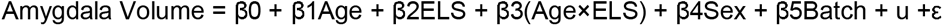

The age × ELS interaction was included to test whether there were volume differences in the amygdala for stressed rats compared to their non-stressed counterparts. The subject ID was included as a random effect to account for repeated measures. To examine potential non-linear trajectory changes in amygdala volume, an additional polynomial mixed-effects model was defined:

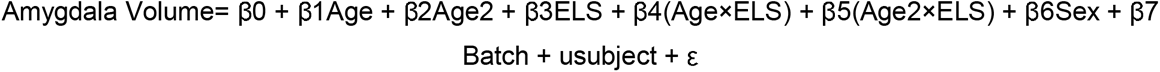

This model included both linear and polynomial age terms, as well as their interactions with ELS. The age² × group interaction was included to test whether the stress group influenced amygdala volume trajectories non-linearly across timepoints. Exploratory analyses examining potential linear and polynomial sex differences were also conducted (Supplementary Note 1; Supplementary Figs 1 and 2).

### 2. Human Adolescent sample

Secondly, we assessed whether childhood trauma was associated with longitudinal trajectories of amygdala volume from adolescence into early adulthood.

#### Data/ Participants

The IMAGEN database contains data collected and processed by the IMAGEN consortium from over 2000 adolescents and their parents. Data have been collected over 10 years across 8 recruitment centres and at 4 successive time points: baseline at age 14 (BL), follow-up 1 at age 16 (FU1), follow-up 2 at age 19 (FU2), and follow-up 3 at age 23 (FU3). A detailed description of recruitment and assessment procedures in the IMAGEN study has been previously reported^56^. Ethical approval was obtained at each of the collection sites (Supplementary Note 2).

No structural MRI scans were collected during the first follow-up visit; therefore, only baseline (BL), follow-up 2 (FU2), and follow-up 3 (FU3) timepoints were included in the analysis. Of the 2,000 adolescents, 1,210 had T1-weighted structural MRI scans at all three timepoints and complete ELS data. Two participants were excluded due to image quality issues. Age data were missing for 18 participants at FU2, 348 participants at FU3, and one participant at both FU2 and FU3. Missing age values were imputed using deterministic methods. For participants with missing FU2 age only, the value was estimated using linear interpolation:

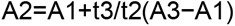

For participants with missing FU3 age only, the value was estimated using linear extrapolation:

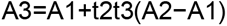

For the participant with missing age at both FU2 and FU3, values were imputed using the median interval approach. As such, the final analysis included 1,208 adolescents (630 females, 578 males) assessed across three timepoints. All siblings were removed from the dataset. The mean age was 14.41 ± 0.39 years at baseline, 19.04 ± 0.74 years at FU2, and 22.91 ± 1.14 years at FU3. Participants were recruited across eight European centres: Paris (n = 172), Nottingham (n = 211), Berlin (n = 96), Dresden (n = 171), Hamburg (n = 135), Mannheim (n = 147), Dublin (n = 112), and London (n = 164).

#### Childhood Trauma Questionnaire

The Childhood Trauma Questionnaire (CTQ) is a 28-item retrospective self-report measure designed to provide an overview of traumatic experiences that may have occurred during childhood and adolescence^57^. There are five subscales, each measuring a different type of traumatic event: emotional abuse, emotional neglect, sexual abuse, physical abuse and physical neglect. Items are scored on a 5-point Likert scale. The entire scale has a Cronbach’s alpha of 0.95^57^. The CTQ was collected at FU2, when the individuals were 19 years old, allowing for a retrospective overview of their experiences throughout childhood and adolescence. CTQ was used as a retrospective measure and therefore represents cumulative exposure before, or concurrent with, the developmental period under investigation. Within our sample, the average CTQ score was 31.74 (±7.95), with scores ranging from 25 to 97.

#### MRI Acquisition Parameters

The scans were acquired on various 3T scanners from eight different sites across Europe, including Siemens (Munich, Germany), Philips (Best, The Netherlands), General Electric (Chalfont St Giles, UK), and Bruker (Ettlingen, Germany). High-resolution, T1-weighted images were obtained using a Magnetisation Prepared Rapid Acquisition Gradient Echo (MPRAGE) sequence, with parameters based on the ADNI protocol (http://www.loni.ucla.edu/ADNI/Cores/index.shtml). All scans had an isotropic voxel size of 1.1mm.

#### Statistical Models

All scans were visually inspected and quality checked. SPM25 was used for the preprocessing following a prevalidated pre-processing pipeline^58^. After inspection, the scans at all timepoints were segmented into probability maps of their tissue classes: grey matter, white matter, CSF, bone and skull. To ensure accurate inter-subject alignment, averaged DARTEL templates were created by aligning the grey matter amongst the images whilst simultaneously aligning the white matter. A flow field for each participant was then generated by comparing the DARTEL template to an average template for each individual. This was then applied to all three time points for each individual to normalise the scans to the standard MNI152 template space whilst conserving the longitudinal consistency. All scans were then smoothed using an 8 mm kernel. Intracranial volume was calculated using the get_totals MATLAB function (http://www.cs.ucl.ac.uk/staff/g.ridgway/vbm/get_totals.m) to adjust brain volumes by head size.

A region-of-interest approach was then taken to identify any volumetric changes in the amygdala. Masks for the left and right amygdala were acquired from NeuroVault (right: https://identifiers.org/neurovault.image:68217, left: https://identifiers.org/neurovault.image:109843) (Gorgolewski et al., 2015). These masks were then combined into one bilateral amygdala mask using FSLmaths (version 6.0.7.14). Values for each voxel within these masks were then extracted for each participant at each timepoint. To assess longitudinal global amygdala volume change associated with ELS, a linear mixed-effects model was used:

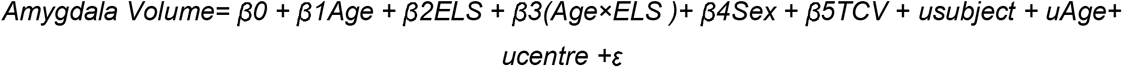

The age × ELS interaction was included to test whether ELS influenced trajectories of amygdala volume across time. Subject and recruitment centre were included as random effects to account for repeated measures and site-related variability. To model individual differences in developmental change, age was included as a random slope at the participant level. Continuous predictors were z-standardised prior to modelling.

To examine potential non-linear trajectory changes in amygdala volume, an additional polynomial mixed-effects model was defined:

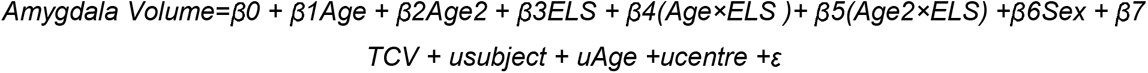

This model included both linear and polynomial age terms, as well as their interactions with ELS. The age² × ELS interaction was included to test whether ELS influenced amygdala volume trajectories non-linearly. Exploratory analyses to examine potential linear and polynomial sex differences in amygdala development associated with ELS were also conducted (Supplementary Note 3).

### 3. Older Adult sample

Thirdly, we tested associations between amygdala volume and ELS in a large-scale mid- to later-life adult sample.

#### 3.1 Total amygdala volume analysis

##### Participants

Data from the UK Biobank (2022 data release), a large-scale biomedical resource that collected health and cognitive data from over 504,000 participants aged 40-69 in Great Britain between 2006 and 2010, were used in this analysis. Ethical approval was granted by the Northwest Multi-Centre Research Ethics Committee (REC reference 11/NW/0382). The current study was approved by the UK Biobank (application number 134918).

Participants were excluded if they had any previous neurological illness or injury, including any disorder affecting the central nervous system, such as demyelinating diseases. Any participants without the required imaging or ELS data were also excluded. The demographics of the resulting sample are displayed in Table 1.

**Table 1.** Demographics of the sample.

|  | Overall | Male | Female |
| --- | --- | --- | --- |
| N | 25,413 | 11,692 | 13,721 |
| Age | 54.85 ($\pm 7.41$) | 55.74 ($\pm 7.49$) | 54.08 ($\pm 7.26$) |
| Ethnicity |  |  |  |
| British | 23,951 | 11,079 | 12,872 |
| Other White Background | 784 | 290 | 494 |
| Asian | 268 | 146 | 122 |
| Mixed Race | 95 | 34 | 61 |
| Caribbean | 70 | 28 | 42 |
| African | 59 | 27 | 32 |
| Other Black Background | 3 | 2 | 1 |
| Other | 119 | 48 | 71 |
| Undisclosed | 64 | 38 | 26 |

##### Materials

To measure ELS, five questions of the ‘Traumatic Events’ questionnaire were selected (Field ID 20489–20491). These questions assess the environment that the participant grew up in and include measures of physical, emotional and sexual abuse as well as physical and emotional neglect. This measure is an English adaptation of the Childhood Trauma Screening^59^, which is based on a shortened version of the CTQ^57^. The CTS strongly correlates with total CTQ scores (r = 0.88; p < .0001) and has strong internal consistency, Cronbach’s alpha = 0.76^59^. Each question was scored on a scale of 0 (‘never true’) to 4 (‘very often true’), with each answer being summed to ascertain a total score resulting in a minimum score of 0 and a maximum score of 20. The average score in this sample was 1.71 ± 2.32. This measure has been widely used to quantify ELS in previous research into childhood stress in the UK Biobank^47^.

Amygdala volume values from the set of Biobank imaging-derived phenotypes were used (Field IDs: 25888-9). Values for each hemisphere were summed to create one value for the region prior to analysis.

##### Acquisition Parameters

Imaging-derived phenotypes generated by the UK Biobank imaging pipeline were used. Detailed acquisition and processing protocols have been described previously^60^. Three imaging centres with identical MR scanners (3.0T Siemens Skyra, software VD13) with Siemens 32-channel receive head coil are used by UKBiobank. To take the T1-weighted scans to IDPs, the UKBiobank normalised the 3D MPRAGE T1w images and then non-linearly warped them to MNI152 using FNIRT (FMRIB’s Nonlinear Image Registration Tool. Tissues are then segmented using FAST (FMRIB’s Automated Segmentation Tool, allowing for grey matter volumes normalised for head size to be generated as IDPs by the UKBiobank team. These analysis pipelines were adapted from the Human Connectome Project^61^.

##### Statistical Analysis

Before the analysis, sensitivity tests were conducted to remove any outliers from the dataset for the amygdala IDP using the same IQR method, where outliers were defined as extreme and removed from the dataset if values exceeded three times the IQR. A total of 5 datapoints were removed. Once the outliers were removed, a linear regression model was conducted to assess the association between amygdala volume and ELS whilst controlling for age and sex:

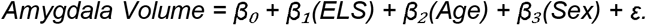

Exploratory analyses to examine potential linear and polynomial sex differences in amygdala development associated with ELS were also conducted (Supplementary Note 4).

#### 3.2 Voxel-Based Morphometry Analysis

We conducted this analysis as a follow-up to the whole-amygdala IDP findings to localise ELS-related effects on a voxel-wise level.

##### Participants

Data from the UK Biobank (2022 data release) were used in this analysis under the same approved Biobank project (application number 134918). A smaller subsample of the previous sample was used to isolate those with the highest incidence of childhood stress, whereas the control sample consisted of individuals who had experienced no childhood stress. Therefore, this sample (n=715) is split into two: high and no ELS. Again, all participants with any neurological illness or injury (including demyelinating disorders) were removed. Any participants without complete T1 images or ELS data were also excluded. Participant characteristics are displayed in Table 2.

**Table 2.**
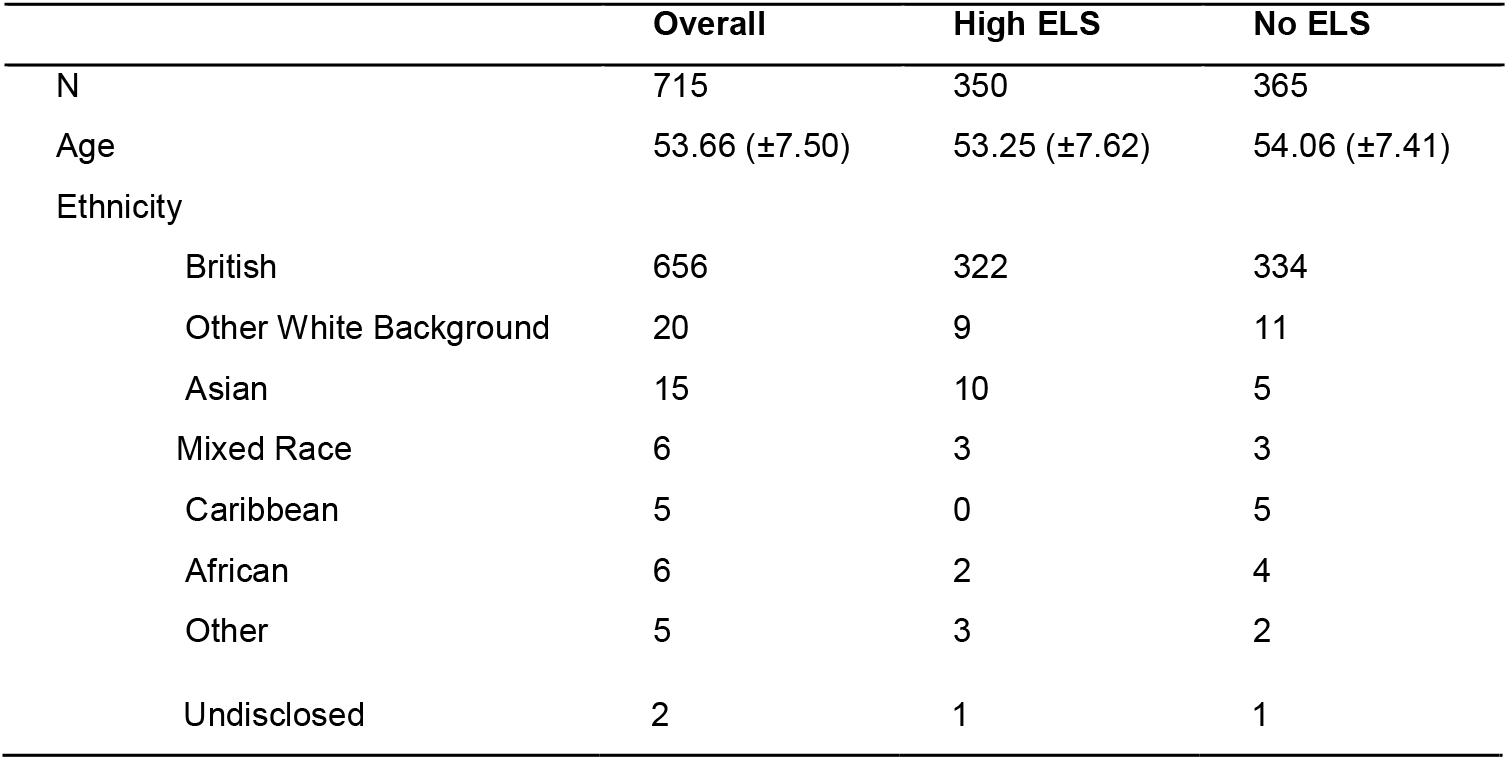
Demographics of the sample for the older adult human VBM analysis.

##### Defining low and high-stress groups

The five-question Traumatic Events questionnaire (Field ID 20489-20491), based on the Childhood Trauma Screening^59^, was used to assess ELS. To maximise sensitivity for voxel-wise analyses and to examine the neuroanatomical correlates of severe adversity exposure, we conducted a follow-up extreme-groups analysis comparing participants reporting high levels of childhood adversity with participants reporting no childhood adversity. As such, we defined two groups: no ELS and high ELS. These groups were defined based on previous work using this measure and defining low- and high-stress margins from the UK Biobank^47^. To define the high-ELS group, we used an a priori threshold of two standard deviations above the sample mean (mean = 5.12, SD = 4.70). Participants with scores greater than or equal to this threshold were assigned to the high-ELS group, whilst participants reporting no childhood adversity (score = 0) formed the no-ELS group. Participants with intermediate adversity scores were excluded from this analysis.

##### Voxel-wise amygdala analysis

All T1w structural MR scans were preprocessed using the same preprocessing pipeline as the IMAGEN analysis^58^. SPM12 was used for this analysis. To target the amygdala, images were masked using the same bilateral amygdala mask created in the previous analysis. Once preprocessing was complete, voxel-wise analyses were conducted using the general linear model. A multiple regression model was defined to examine the relationship between ELS and amygdala volume whilst controlling for age, sex and TCV. At each voxel, the model was defined as:

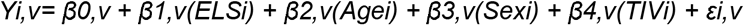

Where Y₍□,□₎ represents the voxelwise grey matter intensity for participant *i* at voxel *v*. Statistical maps were then generated to identify regions in which ELS was significantly associated with amygdala volume. Statistical significance was determined through family-wise error correction within the bilateral amygdala volume implemented through small volume correction in SPM12. Exploratory analyses to examine potential linear and polynomial sex differences in amygdala development associated with ELS were also conducted (Supplementary Note 5).

## Supporting information

All Supplementals

