## Supplementary material for "Early life stress alters lifespan trajectories of amygdala development: a cross-species model of amygdala burnout": All Supplementals

*Supplementary Materials*

**Supplementary Note 1:**

***Rodent sex-difference analysis***

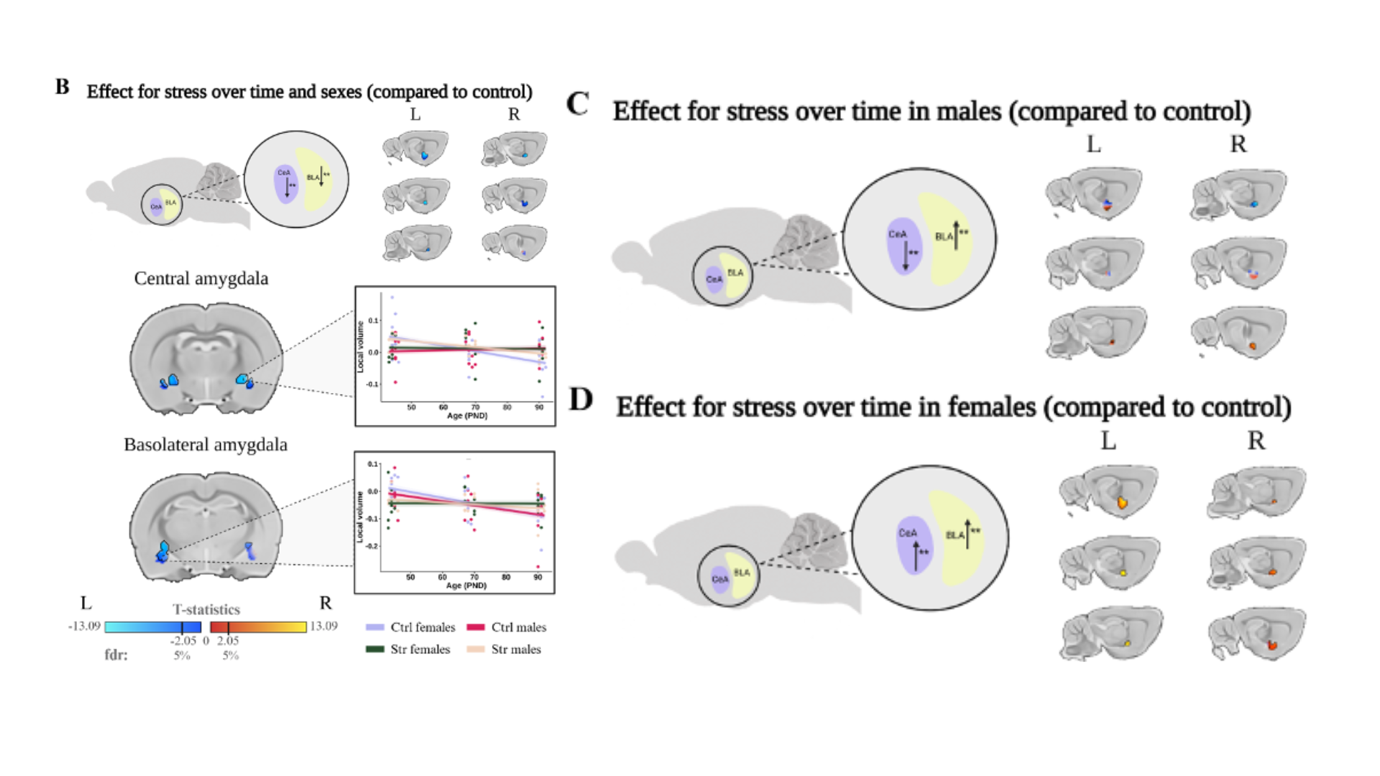
To investigate whether stress-related amygdala changes differed by sex, we tested a three-way interaction (Age × Group × Sex). This analysis revealed a significant interaction (*t* = 2.053, *df* = 871.99, *q* < .05), indicating that the volumetric trajectory of amygdala subregions differed between males and females in response to chronic stress. Sex-stratified analyses revealed only partially overlapping stress-related amygdala patterns. In females, stress exposure was associated with generalised volumetric increases across amygdala subregions with age *(t* = 2.28, *df* = 458.84, *q* < .05). In contrast, males showed a more selective pattern, with stress-related volumetric increases in bilateral BLA and left CeA but decreases in right CeA (*t* = 2.05, *df* = 413.05, *q* < .05).

**A**

**B**

**C**

**Supplementary Fig. 1 Sex-dependent linear effects of chronic restraint stress on amygdala volume.** Structural differences in the amygdala between chronic restraint stress and control groups were examined using voxel-wise models of volumetric trajectories over time. a, Sex-dependent effects identified by the Age × Group × Sex interaction. b, Sex-stratified analysis in males showed volumetric increases in the bilateral BLA and left CeA but decreases in the right CeA. c, Females showed more generalised stress-related volumetric increases in the bilateral BLA and bilateral CeA. Colour maps denote t-statistic maps of group volumetric effects thresholded at a false discovery rate of 5%.

Sex-stratified polynomial analyses revealed less consistent patterns across amygdala subregions, with fewer voxels surviving FDR correction than in the corresponding linear analysis (males: *t* threshold = 2.658, *df* = 455.97, *q* < .05; females: *t* threshold = 2.167, *df* = 411.11, *q* < .05). Although a significant sex interaction was detected, the main stress-related pattern was largely shared between sexes. Both males and females exhibited volume increases within the bilateral BLA and left CeA, with the primary divergence occurring in the right CeA, where females showed increases and males showed decreases.

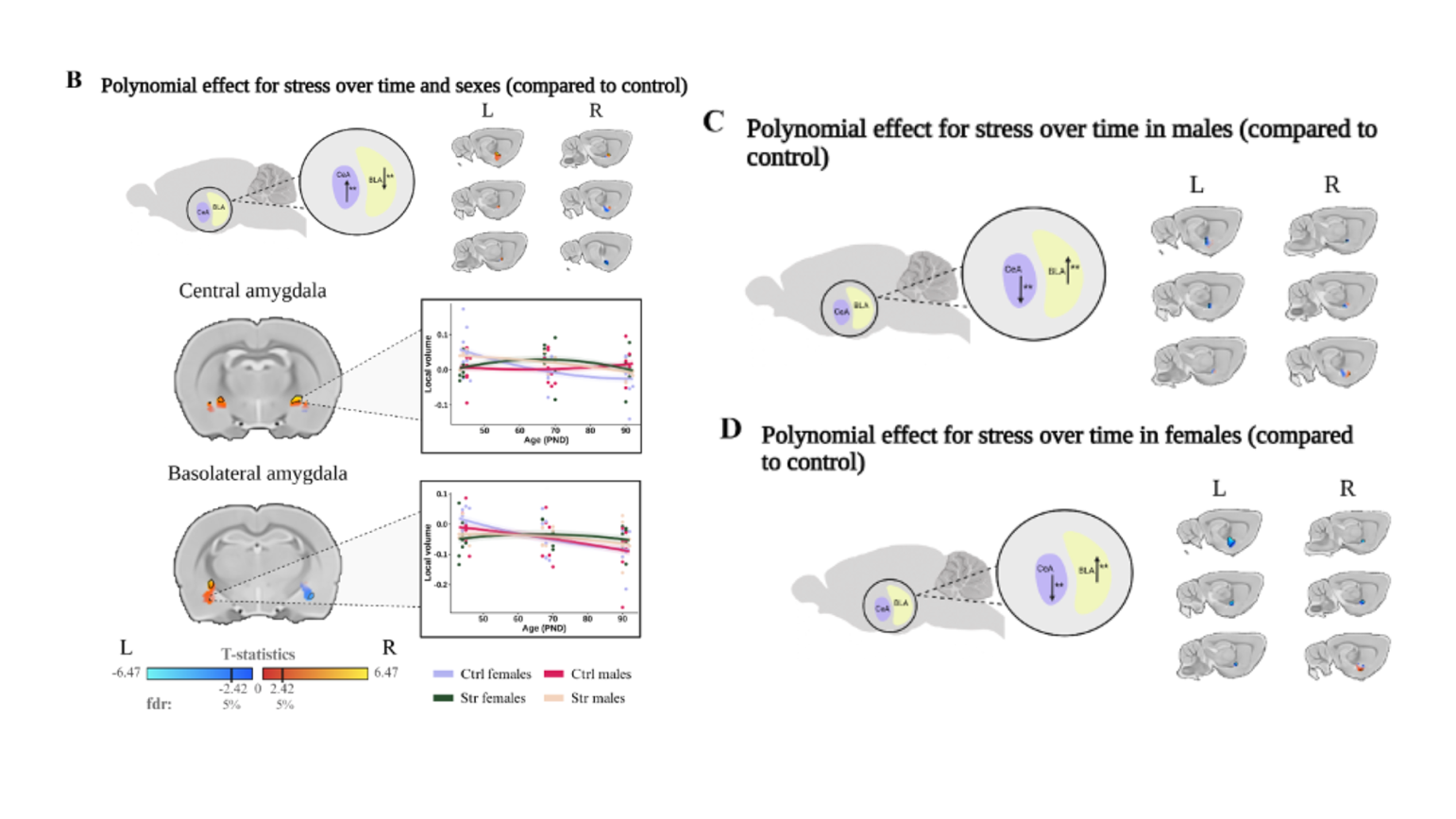

**A**

**B**

**C**

**Supplementary Fig. 2 Sex-dependent polynomial effects of chronic restraint stress on amygdala volume. a**, Sex-dependent effects (Age² × Group × Sex) show divergent patterns between males and females. **b,** Sex-stratified analyses in males revealed selective volumetric increases in right BLA but decreases in left CeA and right CeA. **c,** Females showed more generalised volumetric changes across amygdala subregions, although fewer voxels survived FDR correction than in the corresponding linear analysis. Colour maps denote *t*-statistic maps of group volumetric effects thresholded at False Discovery Rate (FDR) 5%.

**Supplementary Note 2:**

***Ethical approval for the IMAGEN Consortium:***

Ethical approval was granted at each of the data collection sites (London, England: King’s College London Psychiatry, Nursing and Midwifery Research Ethics Subcommittee, King’s College London; Nottingham, England: University of Nottingham Medical School Ethics Committee; Mannheim, Germany: Medizinische Fakultaet Mannheim, Ruprecht Karl Universitaet Heidelberg and Ethik-Kommission II an der Fakultaet fuer Kliniksche Medizin Mannheim; Dresden, Germany: Ethikkommission der Medizinischen Fakultaet Carl Gustav Carus, TU Dresden Medizinische Fakultaet; Hamburg, Germany: Ethics Board, Hamburg Chamber of Physicians; Paris, France: CPP IDF VII (Comité de protection des personnes Ile de France), ID RCB: 2007-A00778-45 September 24, 2007; Dublin, Ireland: TCD School of Psychology REC; and Berlin, Germany: Ethics Committee of the Faculty of Psychology).

**Supplementary Note 3**

***IMAGEN sex-difference analysis***

To investigate whether the association between CTQ score and amygdala volume differed by sex during adolescence, a CTQ × Sex interaction term was included in the linear mixed-effects model. The interaction was not significant (*β* = 0.025, *SE* = 0.212, *t* = 0.119, *p* = .906), providing no evidence that the association between CTQ score and amygdala volume differed by sex.

**Supplementary Note 4**

***UK Biobank IDP sex-difference analysis***

To investigate whether the association between ELS and amygdala grey matter volume differed by sex, an ELS × Sex interaction term was included in the linear regression model. The interaction was not significant (*β* = 3.73, *SE* = 2.74, *t* = 1.36, *p* = .174), providing no evidence that sex moderated the association between ELS and amygdala grey matter volume.

**Supplementary Note 5**

***UK Biobank VBM sex-difference analysis***

To examine whether the association between ELS and amygdala grey matter volume differed by sex, a voxel-wise ELS × Sex interaction was tested using the same second-level multiple regression framework described in the main Methods. No significant voxels were identified for either the positive or negative interaction contrast within the bilateral amygdala following small-volume correction. Thus, there was no evidence that sex moderated the association between ELS and amygdala grey matter volume.

**Supplementary Note 6**

***Sensitivity analysis of random-effects structure.***

The primary analysis modelled participant-specific developmental trajectories by including age as a random slope within participant:

*Amygdala Volume = β0 + β1Age + β2CTQ + β3(Age×CTQ) + β4Sex + β5TCV + usubject + uAge, subject + ucentre + ε*

To assess the robustness of the findings to the specification of the random-effects structure, an additional model including participant-specific random intercepts, but no random age slopes, was fitted:

*Amygdala Volume = β0 + β1Age + β2CTQ + β3(Age×CTQ) + β4Sex + β5TCV + usubject +
ucentre + ε*

The Age x CTQ interaction remained positive and statistically significant in the random-intercept only model, although its estimated magnitude was larger than in the primary random-slope model.

**Supplementary Table 1.**

Robustness of the Age × CTQ interaction across alternative random-effects structures.

| Model | Random-effects structure | β (Age × CTQ) | SE | *t* | *p* |
| --- | --- | --- | --- | --- | --- |
| Primary model | Random intercept + random age slope for participant; random intercept for recruitment centre | 2.38 | 0.96 | 2.49 | .013 |
| Sensitivity model | Random intercept for participant; random intercept for recruitment centre | 5.91 | 0.94 | 6.32 | < .001 |

**Note.** *The primary model provided a significantly better fit than the sensitivity model (likelihood-ratio test: χ²(2) = 342.7, p < .001). Positive Age × CTQ coefficients indicate that higher CTQ scores are associated with steeper age-related associations with amygdala volume. Continuous predictors (age, CTQ and TCV) were scaled to unit standard deviation without mean-centring.*
